## Supplementary Material for "Dimensionality Reduction of Genetic Data using Contrastive Learning"

#### Map Projection

Figure S1 shows the 3D spherical embedding using our contrastive learning framework on the dog dataset, and the resulting 2D embedding from the map projection. Note that the other embeddings have been compressed in the $x$-direction, which might introduce some artificial visual lengthening of the clusters in the $y$-direction.

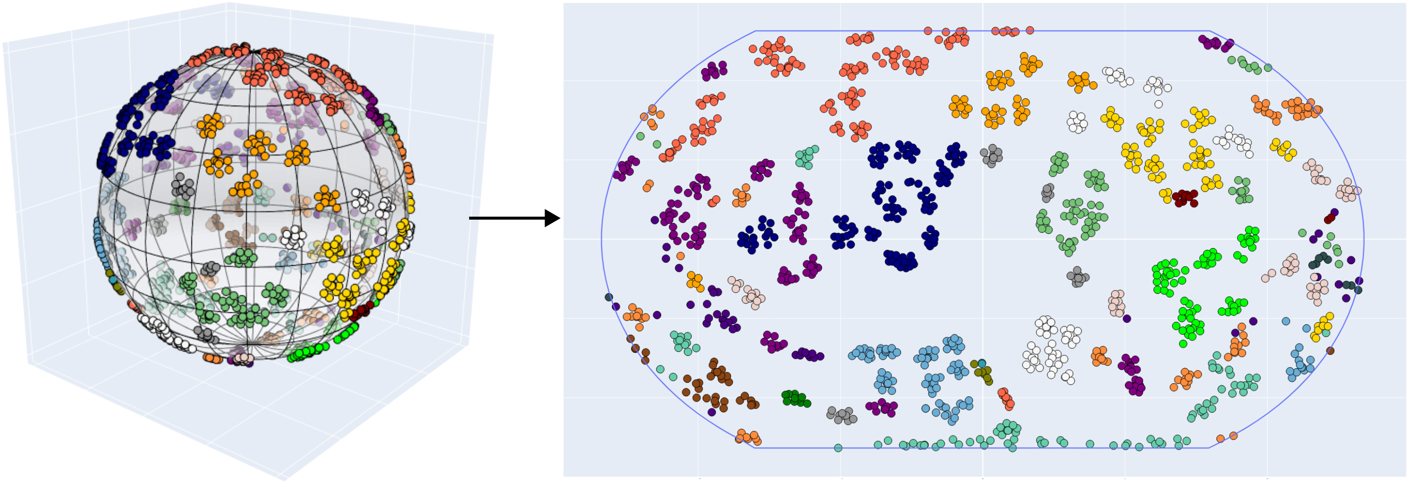

Figure S1: Illustration of the spherical embedding and the resulting 2D embedding using the map projection of the dog dataset after having rotated the sphere.

When applying the map transformation, it may matter how the 3D coordinate system is defined. The orientation of the sphere affects which samples will be mapped closer to the boundary than others. By perturbing the latitude and longitude we can obtain different embeddings. This is done by iteratively perturbing either the latitude or longitude by $2\pi/10$ at a time, until the sphere has rotated once along both the $x$ and the $z$ axes. Figure S2 shows in the left plot how this perturbation affects the accuracy of KNN classification on the projection for masked Human Origins data (the same 3D embedding as Figure 8 in the main manuscript), and the right plot shows the projection with the worst perturbation. Here both the East and Central/South Asia are split to opposite sides of the embedding which results in a visually poor embedding. Here the KNN score should be seen as a proxy for the perceived quality of the 2D visualization changes with the orientation of the sphere. Note that it only changes a couple of percentage points, and in the main text we use the KNN score on the spherical embedding to evaluate performance consistently.

#### Spherical UMAP embeddings

In the UMAP implementation, one can train the embedding using the haversine distance, which outputs latitude and longitude, which can be translated into an embedding on the unit sphere. To give an apples-to-apples comparison to our 3D spherical embeddings, we do the comparison here on the dog dataset.

As in the main text, we look at two values for the number of neighbors, 3 and 30. Table S1 compares the result of the 3D spherical UMAP embeddings to plain 2D UMAPs. We see worsened results across the board using the spherical embeddings.

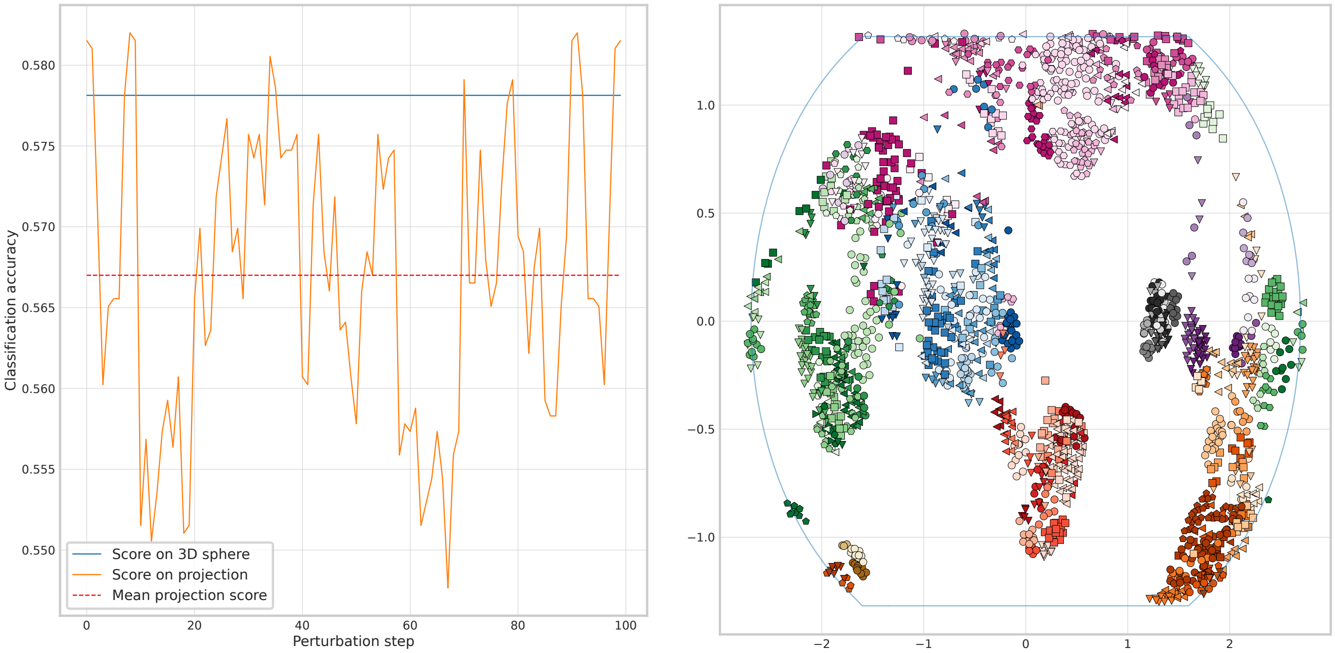


Figure S2: Left: KNN classification accuracy for different perturbations of latitude and longitude of the contrastive learning embedding. The blue line is the KNN accuracy on the 3D embedding, and the dashed red is the mean of the resulting 2D projections. Right: The 2D projection with the perturbation which resulted in the worst KNN classification accuracy.

Table S1: Performance comparison of UMAP 2D embeddings with UMAP embeddings on the unit sphere:

| **Method** | Neighbors | **Glob. score(G)** | **Subpop acc.(L)** | **Validation (GE)** |
|---|---|---|---|---|
| UMAP 2D | 3 | 0.3107 | 0.9207 | 0.9114 |
| UMAP 2D | 30 | 0.4294 | 0.9022 | 0.8229 |
| UMAP Spherical | 3 | 0.1525 | 0.7039 | 0.5055 |
| UMAP Spherical | 30 | 0.3277 | 0.8395 | 0.4908 |

### 3-Dimensional Embeddings and Map Projection for PCA and t-SNE

Here we show the resulting embeddings of the dog dataset when we use the same methodology that we use for our contrastive learning implementation. First, 3D embeddings were created using PCA and t-SNE, the coordinates were normalized to the sphere, and the Equal Earth map projection was applied. Figure S3 shows the embeddings, and Table S2 compares the performance in terms of population classification accuracy. We see that for both PCA and t-SNE, this worsens the embedding quality. This is most likely due to the 3D embeddings utilizing the full three-dimensional space, and then collapsing to the sphere. This in comparison to our method, where it is trained directly on the sphere.

This shows that computing 3D embeddings and using a map projection is not necessarily the best option out of the box, as it produces worse embeddings for PCA and t-SNE. It is, however, something we can exploit in the deep learning setting, to be able to train in 3D while visualizing in 2D. For t-SNE we have used the same set-up as in the main article, with PCA-preprocessing, and using a perplexity of 30.

Table S2: Classification performance of the different methods on the dog dataset using a KNN classifier with $k = 3$ on the embedding coordinates.

| **Method** | **Superpop clust.(G)** | **Subpop acc. (L)** | **Validation (GE)** |
|---|---|---|---|
| t-SNE 2D | 0.5480 | 0.9271 | 0.9041 |
| t-SNE 3D + projection | 0.3955 | 0.8598 | 0.8708 |
| PCA 2D | 0.4011 | 0.3653 | 0.3210 |
| PCA 3D + projection | 0.4633 | 0.3081 | 0.3173 |

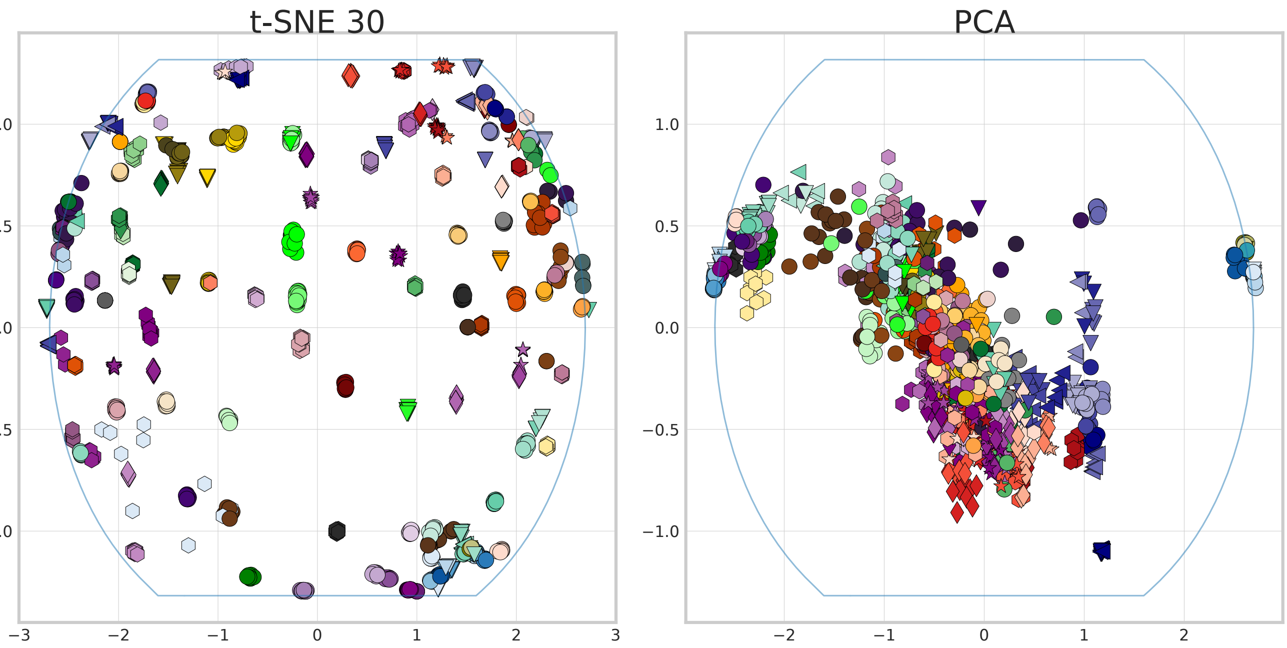


Figure S3: Resulting embeddings t-SNE and PCA when following the same procedure as we do for our contrastive learning implementation. Here we compute 3-dimensional PCA and t-SNE embeddings, normalize to the sphere, and use a map projection.

#### Placement of Centroid

Figure S4 shows where the centroid is placed, depending on the weighting of the negative sample. We argue that the bottom plot, where the negative coordinate is weighted twice, yields a more natural centroid placement. This heuristic is mostly based on the fact that the anchor and the positive are two views of the same sample.

#### Embeddings of Simulated Data

In the main manuscript we focus on real empirical data. To confirm that the contrastive learning model performs reasonably, we also here show embeddings of four cases of simulated data for all embedding methods that we consider. For the experiments in this section we have used msprime version 1.3.3 [1] to set up the simulations and create all datasets. The simulated populations are inspired by the scenarios previously considered for demonstrating possible pitfalls in using PCA [2].

In our first scenario, we consider two populations (pop1 and pop2) that split from an ancestor population of size $N = 10000$, $N/2$ generations ago, and a third population pop3 consisting of a recent 80/20% admixture of pop1 and pop2 5 generations ago. From this, we generate 500 individuals from pop1, and 250 from both pop2 and pop3. This way, we have an imbalanced dataset with respect to the sample sizes and a known admixture pattern. We used a sequence length of $10^7$, a recombination rate of $10^{-8}$, and a mutation rate of $10^{-8}$, which resulted in a dataset of 1000 individuals and a SNP count of 32993. Embeddings of this dataset are shown in the leftmost column in Figure S5.

The three other datasets are variations of a linear stepping stone model, where populations only have genetic exchange with their closest neighbor. We consider 10 populations, with $N = 1000$ in each, and an effective migration rate of $Nm = 1$. In the first case, we simulate 100 individuals from each population. In the second case, we use the same dataset, but only consider samples from populations 1-4 and 7-8. This means that we should expect a gap between populations 4 and 7. As a last dataset, we also allow migration between populations 1 and 10, creating a ring. This results in three different datasets, which we denote *Stepping Stone*, *Stepping Stone Clustered*, and *Stepping Stone Ring*. The embeddings are shown in Figure S5 columns 2, 3, and 4, respectively. We use the same settings for sequence length and recombination and mutation rates as in the admixture dataset. The three datasets have (1000, 37880), (600, 37880), (1000, 36098) individuals and non-fixed SNPs, respectively.

The purpose of this exploration was to verify that our method performs reasonably, and that there are no surprising artifacts on simulated cases where we know the population structure. Therefore, we focus on the qualitative differences and similarities of the embeddings. In our contrastive embeddings, we have rotated the coordinate system before applying the map projection to give a visualization that is easier to interpret. The UMAP and t-SNE embedding have been created

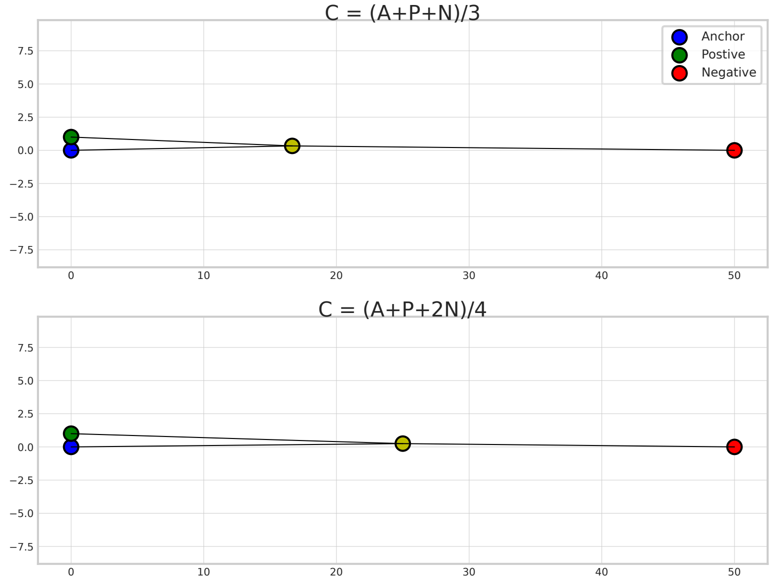


Figure S4: Position of the centroid based on different scaling of the negatives.

in the same manner as in the main manuscript, by applying the methods on the first 100 PCA components of the data. We have used a perplexity and number of neighbors of 100, for the methods to perform better on the stepping stone datasets.

Overall, our method produces reasonable embeddings throughout. For example, compared to UMAP, our contrastive method produces an embedding that does “close the circle” in the ring dataset when one considers the wrapping property inherent in our embedding. Unlike popvae, our contrastive embedding shows a distinct gap between pop4 and pop7 in the clustered stepping stone dataset. Our method also shows a more apparent linear relationship between the populations in the stepping stone embeddings, where popvae sometimes stacks the populations, like pop4 and pop5. The popvae embeddings generally stand out, since they push samples from the origin creating distorted embeddings of especially pop1 and pop10 in the stepping stone datasets. This behavior is also seen in the empirical datasets shown in the manuscript.

The UMAP and to some extent popvae embeddings of the admixture dataset seem to be able to best visualize that pop1 was more represented in the admixture event. This is not something that our method shows, but we do not claim that our embedding coordinates correlate directly to genetic distance, but only that the embedding is topologically correct given the dataset. All methods except for PCA and our proposed method create visually distinct gaps between populations in the Stepping Stone and Stepping Stone Ring scenarios, with a few individuals ending up in a cluster different than their origin population. This implies that there are some individuals with a short distance if one would compare the original genotype vectors, that still end up with an apparent gap in the resulting embedding. This effect is similar to the overclustering effect present in the All of Us visualizations that we discuss in the main text.

### References

[1] Franz Baumdicker, Gertjan Bisschop, Daniel Goldstein, Graham Gower, Aaron P Ragsdale, Georgia Tsambos, Sha Zhu, Bjarki Eldon, E Castedo Ellerman, Jared G Galloway, Ariella L Gladstein, Gregor Gorjanc, Bing Guo, Ben Jeffery, Warren W Kretzschumar, Konrad Lohse, Michael Matschiner, Dominic Nelson, Nathaniel S Pope, Consuelo D Quinto-Cortés, Murillo F Rodrigues, Kumar Saunack, Thibaut Sellinger, Kevin Thornton, Hugo van Kemenade, Anthony W Wohns, Yan Wong, Simon Gravel, Andrew D Kern, Jere Koskela, Peter L Ralph, and Jerome Kelleher. Efficient ancestry and mutation simulation with msprime 1.0. *Genetics*, 220(3):iyab229, 2022.

[2] John Novembre and Matthew Stephens. Interpreting principal component analyses of spatial population genetic variation. *Nature Genetics*, 40(5):646–649, May 2008.

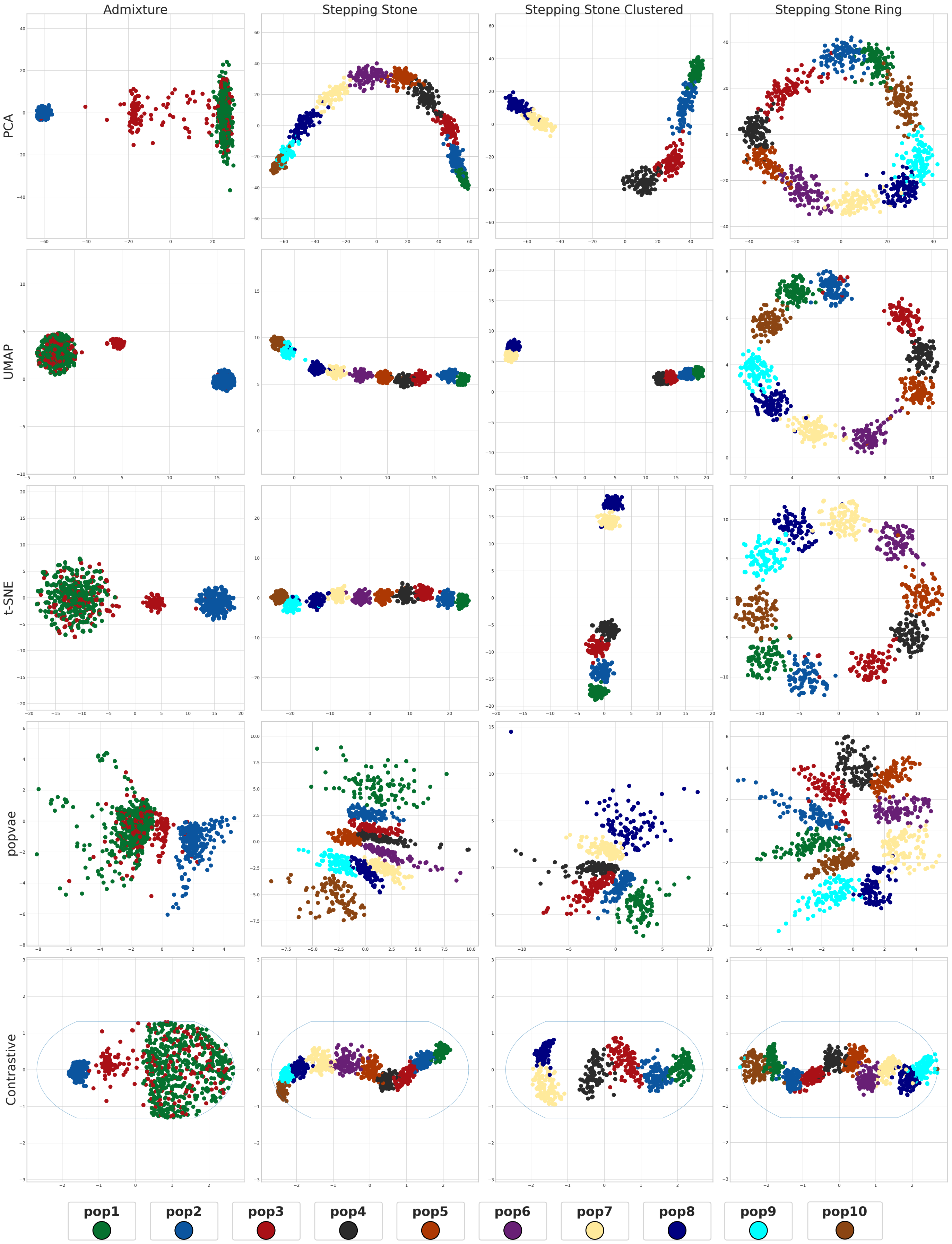


Figure S5: Embeddings of four cases of simulated data, using PCA, UMAP, t-SNE, popvae, and our contrastive method.
